# Prediction of Fine-tuned Promoter Activity from DNA Sequence

**DOI:** 10.1101/030049

**Authors:** Geoffrey H. Siwo, Andrew K. Rider, Asako Tan, Richard S. Pinapati, Scott Emrich, Nitesh Chawla, Michael T. Ferdig

## Abstract

The quantitative prediction of transcriptional activity of genes using promoter sequence is fundamental to the engineering of biological systems for industrial purposes and understanding the natural variation in gene expression. To catalyze the development of new algorithms for this purpose, the Dialogue on Reverse Engineering Assessment and Methods (DREAM) organized a community challenge seeking predictive models of promoter activity given normalized promoter activity data for 90 ribosomal protein promoters driving expression of a fluorescent reporter gene. By developing an unbiased modeling approach that performs an iterative search for predictive DNA sequence features using the frequencies of various k-mers, inferred DNA mechanical properties and spatial positions of promoter sequences, we achieved the best performer status in this challenge. The specific predictive features used in the model included the frequency of the nucleotide G, the length of polymeric tracts of T and TA, the frequencies of 6 distinct trinucleotides and 12 tetranucleotides, and the predicted protein deformability of the DNA sequence. Our method accurately predicted the activity of 20 natural variants of ribosomal protein promoters (Spearman correlation *r* = 0.73) as compared to 33 laboratory-mutated variants of the promoters (*r* = 0.57) in a test set that was hidden from participants. Notably, our model differed substantially from the rest in 2 main ways: i) it did not explicitly utilize transcription factor binding information implying that subtle DNA sequence features are highly associated with gene expression, and ii) it was entirely based on features extracted exclusively from the 100 bp region upstream from the translational start site demonstrating that this region encodes much of the overall promoter activity. The findings from this study have important implications for the engineering of predictable gene expression systems and the evolution of gene expression in naturally occurring biological systems.

**Author Summary:** Gene expression is the first step at which information encoded in DNA is transcribed into RNA. Predicting gene expression from DNA sequence can provide insights into the natural variation of gene expression underlying various phenotypes and direct the engineering of genes of desired activity, for example in industrial processes. While several studies show that gene expression is influenced by DNA sequence. its quantitative prediction from DNA sequence alone remains a challenging problem. Unfortunately, studies aimed at developing quantitative models for gene expression prediction are not directly comparable because most have used distinct data sets for training and evaluation. and many of the methods have not been independently verified. Open innovation challenges in which a problem is posed to a wide community provide a framework for independent verification of the performance of various computational methods using the same benchmark data sets and statistical procedures. Here. we describe the best performing computational model amongst those of 20 other teams in the DREAM6 Gene Expression Prediction challenge. We show that a highly predictive gene expression model can be obtained by an unbiased. data-driven approach that makes little assumption on the role of known mechanisms for gene regulation.

## Introduction

Transcription is a fundamental step in the decoding of information encoded in DNA into phenotypes. Therefore. knowledge of transcriptional regulation is crucial for understanding the natural variation of gene expression [1-5] and for the accurate engineering of predictable gene expression systems [6-8]. While transcriptional regulation is one of the most highly studied areas in biology the ability to quantitatively predict gene expression from DNA sequence remains inadequate [9,10]. Knowledge of transcription factors and their cognate binding sites continues to grow and has enhanced our ability to make qualitative predictions about gene expression. For example. a number of transcription factors are now well known to be involved in differentiation of stem cells into specific cell types leading to potentially clinically useful applications such as induced pluripotent stem cells [11]. In spite of this progress, only limited quantitative predictions of gene expression are possible [6-8,12,13]. Knowledge that promoter sequences of genes encode both qualitative (e.g. when to switch a gene on and off) and quantitative properties (e.g. precise levels and noise) of gene expression is implied by the heritable nature of these attributes [1-3,14]. It is becoming increasingly clear that while transcription factors are critical in gene regulation. regulatory outputs are ultimately determined by co-operation between regulators in complex circuits [15-17] and with chromatin states [18-21]. In particular, transcription factors compete for DNA binding sites with nucleosomes [22,22,23]. The information for nucleosome binding is largely encoded in the DNA sequence [24-27]. even though *in vivo* nucleosome occupancy is highly dynamic [25,28,29]. Quantitative models of gene expression, therefore, benefit from the integration of nucleosome and transcription factor binding data [10,23,30].

A key barrier to quantitative modeling of gene expression using promoter sequence has been the lack of experimental methods for accurately measuring transcript levels. DNA microarrays and RNA-seq are the most widely-used systems for measuring transcript abundance, but this measurement can reflect many effects including promoter sequence, genomic position of a gene and post-transcriptional regulation of mRNA levels by processes like mRNA degradation. In addition, microarray and RNA-seq can be affected by systematic biases arising from sequence dependent hybridization kinetics [31] and sequence dependent read-depth coverage [32], respectively. To overcome these limitations, approaches based on promoters fused to fluorescent reporters have been developed to generate direct, real-time measurement of promoter activity with high accuracy [33]. This has been applied in large libraries of synthetic bacterial promoters thereby generating new insights on combinatorial cis-regulation [8]. It was not until recently that the first large-scale library of naturally occurring promoters of any eukaryote fused to yellow fluorescent protein (YFP) became available [30]. 110 yeast ribosomal protein (RP) promoters were fused to YFP and integrated into a different strain at a fixed genomic location, hence alleviating both post-translational and genomic context related effects [30]. Consequently, this data set is very well poised for the computational modeling of the relationship between promoter sequence and transcription activity of a eukaryotic promoter.

To provide a fair assessment of the relationship between promoter sequence and quantitative transcript levels, the Dialogue for Reverse Engineering Assessments and Methods (DREAM) organized an open community challenge in 2011 (details of the challenge as well as an overview of participating teams is provided in reference [34]), inviting participants to address this question using promoter activities of the RP promoter library that was not yet published [30]. Participants were provided with the activities of 90 promoters and their corresponding promoter sequences and challenged to predict the activity of 53 promoters whose activities were known only to the organizers of the challenge (Figure 1A). After a period of three months, the challenge organizers independently assessed the performance of models from 21 teams using four different statistical tests. Our team, FIrST, attained the best performance status on the basis of a combined score by the DREAM consortium in predicting the activities of these 53 promoters (Spearman correlation between predicted and actual activities = 0.65, *P* = 0.002). Our approach was built upon three key propositions: i) transcription factor binding and nucleosome binding, as well as other regulatory signals are encoded in DNA [9,10,12,27]’ ii) if i) is true, then explicit prior knowledge of transcription factor and nucleosome binding is not a mandatory pre-requisite for prediction of promoter activity if training data is available. That is, an unbiased approach that explores the associations between DNA sequence patterns and promoter activity should be able to rediscover patterns that relate to the observed activity. To do this, we used machine learning methods to iteratively explore the association between promoter activity and DNA sequence patterns in 100 bp windows of promoter sequence. We considered sequence patterns such as k-mers (k = 1 to k = 5), homopolymer stretches, nucleosome binding and three mechanical properties of DNA (bendability [35], deformability [36] and stiffness [37]). Based on iterative exploration of different machine learning models, we established that a support vector machine (SVM) was the most predictive of promoter activity based on specific sequence patterns in the 100 bp upstream of the translation start site (TrSS). Our model outperformed those which applied transcription factor binding sites of known RP promoters [34], implying that other sequence patterns besides transcription factor binding sites can help in fine-tuning gene expression. Indeed, among the predictive features employed by our model were poly(dT-dA) tracts that occlude nucleosomes; these have since been applied to fine-tune gene expression beyond resolutions attainable by transcription factor site mutations [38]. Our study expands the understanding of sequence patterns that could potentially be useful in engineering fine-tuned gene expression.

**Fig. 1.**
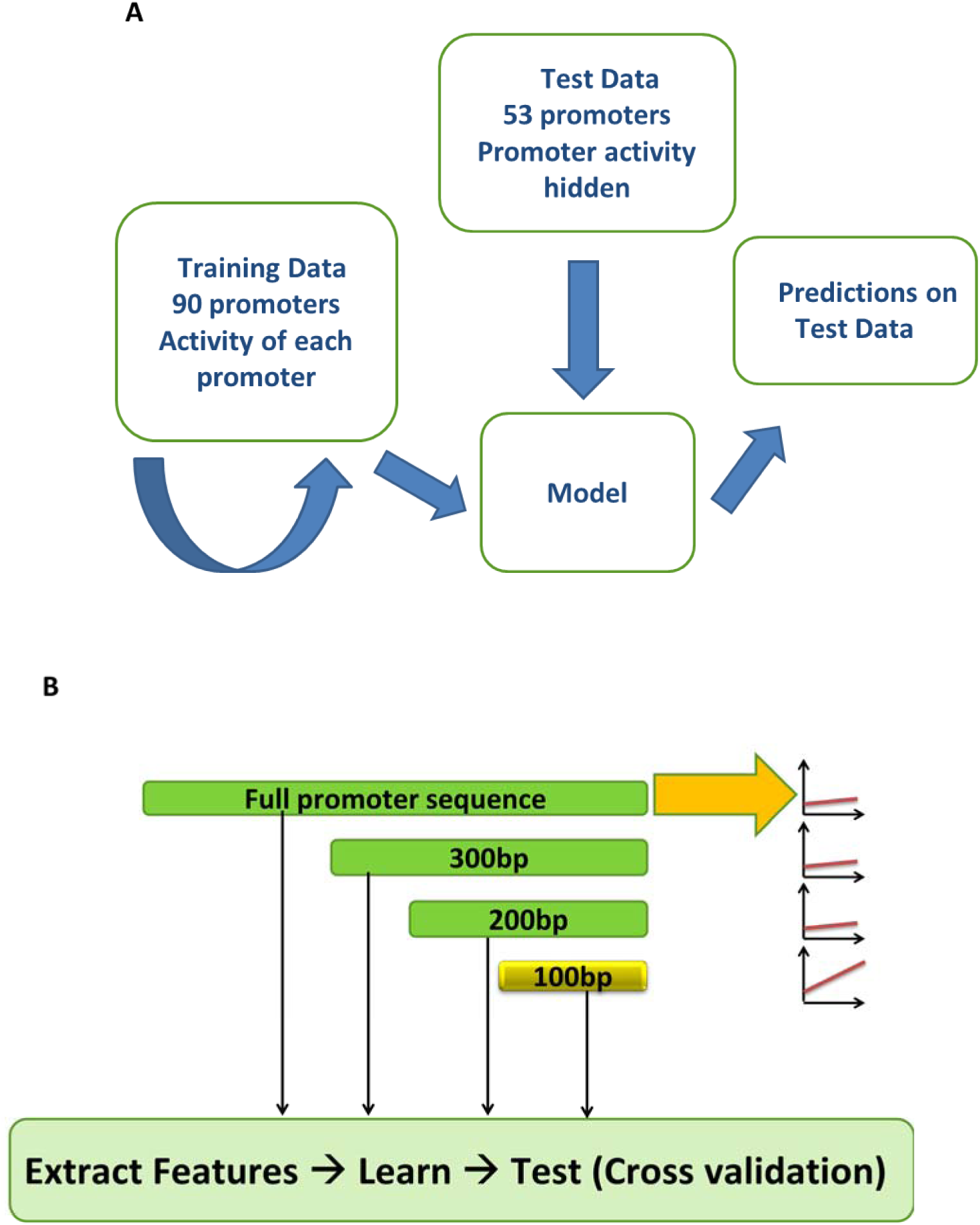
Summary of the DREAM6 gene expression challenge. (A) Training data consisted of DNA sequences for 90 yeast RP promoters whose activities were experimentally determined [30,34]. DNA sequences for blinded test set of 53 promoters whose activity was hidden also experimentally determined but withheld from the challenge participants was also provided. (B) Outline for strategy of modeling promoter activity. Each promoter was segmented into 100 bp non-overlapping windows with the full promoter regarded as a separate window. For each window, DNA sequence features were extracted and feature selection using a linear regression wrapper performed prior to machine learning. Performance of machine learning models trained on each window was determined in 5- and 10-fold cross-validations using Pearson correlation.

## Results

### Promoter Activity is Highly Predictable using the 100 bp Upstream Region from TrSS

The challenge organizers provided DNA sequences and promoter activities-the average rate of YFP production from each promoter, per cell per second, during the exponential phase - for 90 RP promoters (training set) and another set of 53 promoters whose activity was withheld from participants (test set) [30]. We first partitioned the promoter sequences into 100 bp non-overlapping windows, extracted specific DNA features from each window and considered the full promoter sequence as its own window (Fig. 1B). The features considered were k-mers (k = 1 to 5), length of homopolymeric stretches, nucleosome positioning and DNA mechanical properties (bendability, deformability and stiffness). For each window, we performed feature selection using a linear regression wrapper, then explored three different machine learning methods (SVM, linear regression and regression trees) to learn the association between features in the window and promoter activity (Fig. 1B). The performance in each window was assessed by Pearson correlation using 5- and 10-fold cross-validations on the training data. We observed very poor correlation (r << 0.5) between predicted and actual promoter activities except when using the window comprising 100 bp from the TrSS. Therefore, we focused the SVM model on this window using 23 features (Table 1) selected by the linear regression wrapper. A test of this model on 1000 randomized splits of the data (66% training and 34% testing sets) gave an average Pearson correlation of 0.78. The performance of machine learning models can be biased by the training/ test data set used. Therefore, to reduce this bias, we obtained an additional 500 SVM models trained on randomly sampled sets of 80% of the data and validated on the remaining 20%. In the DREAM test set (activities for this set were withheld from participants), we used the SVM models to make predictions for each promoter. For each promoter, the predicted activity was the average of predictions across all the ensemble of SVMs based only on the 100 bp upstream of the TrSS. These predicted activities were then submitted to the DREAM consortium for evaluation [34].

**Table 1.** DNA sequence features predictive of promoter activity.

| DNA Feature | Description |
| --- | --- |
| Mononucleotides | Frequency of G |
| Dinucleotides | Frequency of GT |
| Trinucleotides | Frequency of 6 trinucleotides |
| Tetranucleotides | Frequency of 12 tetranucleotides |
| T-tracts | Length of T-tracts |
| TA-tracts | Length of TA-tracts |
| DNA deformability | Negatively correlated to activity |

A total of 21 teams participated in the challenge (https://www.synapse.org/#!Synapse:syn2820426/wiki/). Predictions from our team had a Spearman correlation of 0.65 (*P* = 0.002, Fig. 2A) to the actual activities, Pearson correlation of 0.65 (*P* = 0.003), chi-squared χ2 distance metric of 52.62 (*P* = 0.508) and R^2^ statistic measuring the difference in ranks between predicted and actual promoter activities of 35.85 (*P* = 0.004). The *P*-values were generated from the probability of obtaining a comparable or lower performance using a null distribution in which predictions were made by randomly choosing an activity for each promoter amongst all the 21 participating teams. A combined score based on the negative logarithm (base 10) of the geometric mean of the *P*-values for all the 4 scores ranked our team first [34] (Fig. 2B), with significant *P*-value in three out of four of the statistical tests used for evaluation. Further, although we were not ranked first in the χ2 distance metric, our model performed the most consistently across the multiple assessment metrics, suggesting a robustness of the method. A detailed comparison of the teams was published previously by the DREAM consortium [34].

**Fig. 2:**
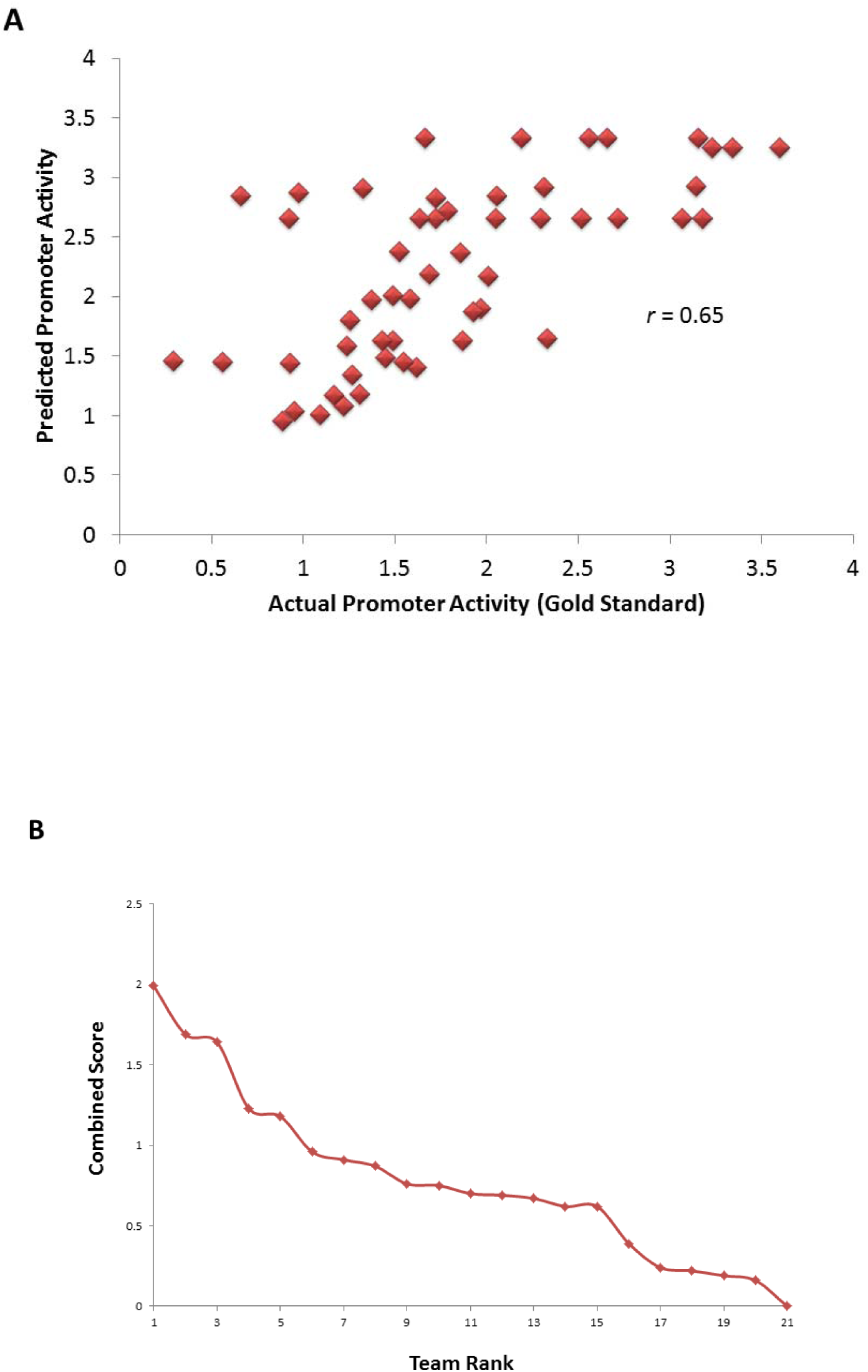
Performance of the SVM model on validation test set by the DREAM consortium. (A) Correlation between predicted activity by the SVM model and actual promoter activity of 53 promoters whose activity was not available to participants. (B) Performance of team FIrST relative to other 20 teams based on a combined score.

### Biological Significance of selected Features

The final SVM models utilized only 23 features consisting of the frequencies of the mononucleotide G, dinucleotide GT, 6 different trinucleotides, 12 different tetranucleotides, length of poly(dT) and poly(dA-dT) tracts (Table 1). The relative importance of these features based on weights for the SVM models is provided in the Supplementary Material. The feature with the highest weight was the frequency of the mononucleotide G, correlating negatively with promoter activity. For many of these features there was no clear link to underlying mechanisms of gene regulation. However, it is possible that some of the k-mers may be implicitly linked to transcription factor binding sites. That is, the combination of different k-mer features could capture the binding motifs of specific transcription factors. For example the second most important feature in the SVM was the tetranucleotide ACCC which also occurs in the *Rap1* binding site motif [39]. In addition, frequencies of different k-mers could impact the DNA mechanical structure [40]. Among the features identified by the SVM model were poly(dT) and poly(dT-dA) tracts which influence the rigidity of DNA [24,26], thereby directly impacting nucleosome binding. Furthermore, insertion of poly(dT-dA) sequences into promoters can be used to regulate gene expression to a finer degree and at more gradual intervals than could be attained by transcription factor binding site mutations [38]. Some transcription factors are also highly dependent on the ability of DNA to bend [41-43]. In particular, TATA binding protein (TBP), which binds to the TATA box, is important for regulating the activity of RP promoters [42,44,45]. Another directly biologically relevant feature identified by the SVM was the deformability of DNA [36,46]. Promoters of low activity had more deformable DNA than those of high activity (Fig. 3, *P* = 0.008). This was particularly evident at 40 to 60 bp from the TrSS when comparing the top 20 promoters with the highest versus those with the lowest activity (Fig. 3).

**Fig. 3:**
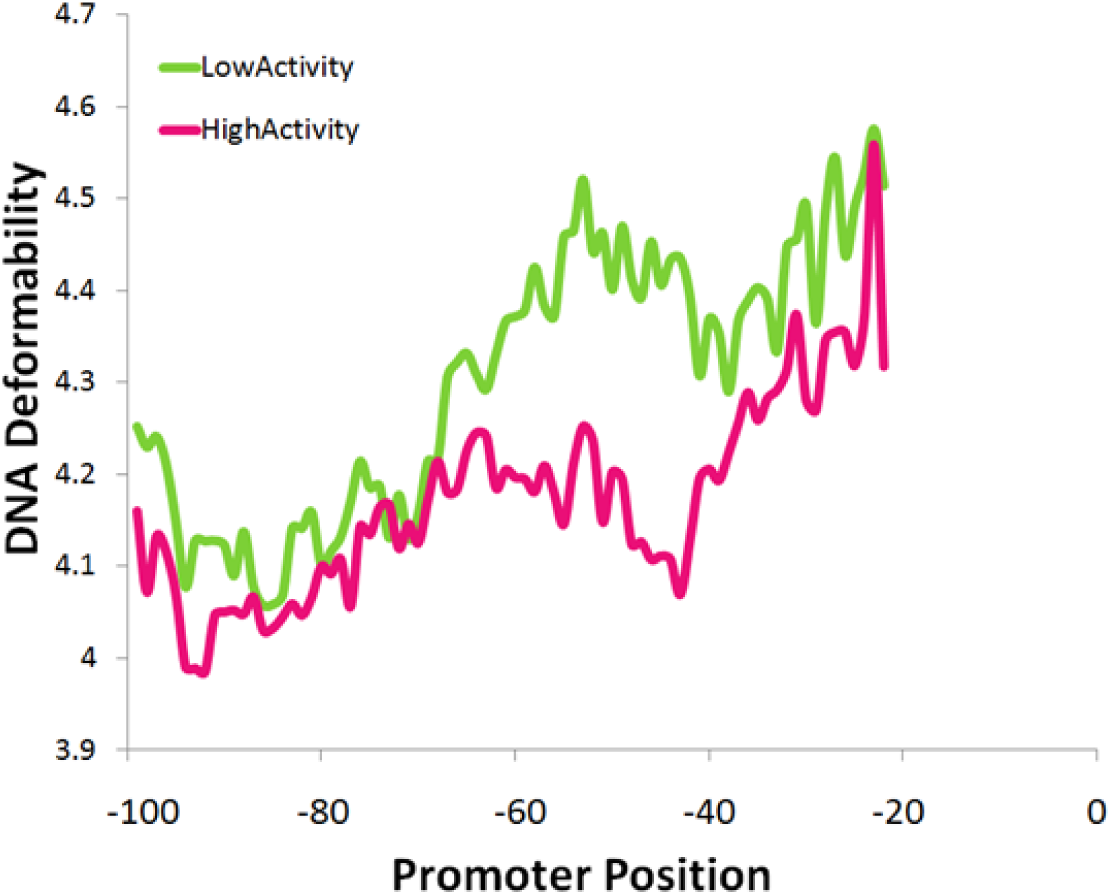
Relationship between protein deformability of promoters and activity. Among the top 20 promoters with extreme activities (high and low), significant deviation in deformability occurs at the -40 to -60 bp region from the TrSS (T-test *P* = 0.008).

Finally, some of the features may affect mRNA stability, especially given their potential location downstream of the transcription start sites (TSS). Besides sequence features in the 5’UTR that are close to the TSS could affect transcription, translation and mRNA stability.

### Error Profile of SVM Promoter Activity Model

Understanding the biases in prediction accuracy could provide biological insights into promoter classes and allow for refinement of models. Therefore, we investigated relationships between the nature of the test promoters and the magnitude of prediction error made by our model. Among the 53 test promoters provided by the DREAM challenge, 20 were natural yeast RP promoters while 33 were variants of these promoters with specific synthetic mutations introduced. These mutations included changes in the binding sites of the TBP, *Rap1, Fhl* and *Sfp1,* as well as introduction of nucleosome disfavoring sequences and random mutations. At the time of the challenge, participants were not aware of these mutations. The performance of our model on the set of natural promoters was much higher (Pearson correlation = 0.73, *P* = 0.0003) compared to that for the mutated promoters (Pearson correlation = 0.57, *P* = 0.0005). The prediction error was significantly less for natural promoters versus the mutated promoters (Student’s t-test, *P* = 0.01, Fig. 4A). This could partly be due to the composition of the training set, which contained only natural promoters. Similar poor performance was also observed in the models obtained from other teams [34]. In addition, most of the synthetic mutations were introduced at promoter locations residing outside of the 100 bp region from the TrSS and could not therefore be detected by our model. We also examined the correlation between the observed promoter activity and the prediction error. Promoters of low activity had larger prediction error (Pearson correlation between promoter activity and prediction error = -0.31, *P* = 0.02, Fig. 4B). Notably, natural promoters had slightly lower activity compared to synthetic promoters (*P* = 0.02) so the correlation between activity and prediction error may be a consequence of the low predictability of synthetic promoters. Thus, future models may benefit from data on activities of mutated promoters, which could enable a more accurate modeling of the impact of mutation on specific transcription factor binding sites.

**Fig. 4:**
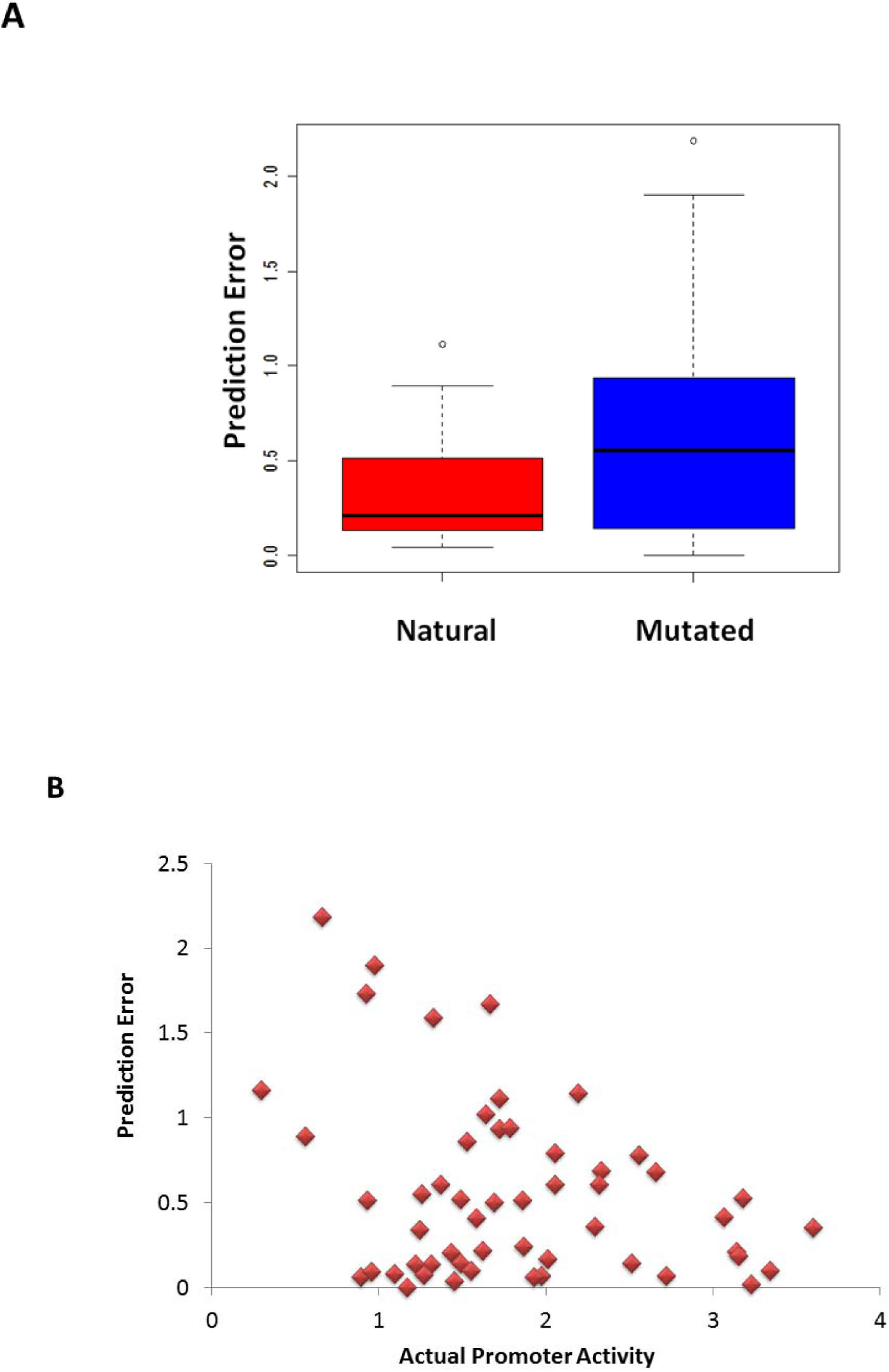
Dependence of prediction error on promoter class or activity. (A) Natural promoters had a lower prediction error compared to synthetically mutated promoters. (B) Prediction error is negatively correlated to promoter activity.

## Discussion

The quantitative modeling of gene expression has the potential to enhance our understanding of how gene regulation is fine-tuned in natural populations and has implications for the design of predictable gene expression systems. The DREAM6 challenge dataset for promoter activity prediction was a unique opportunity to evaluate the predictability of gene expression from its promoter sequence. Given that all promoters were derived from natural yeast RP promoters that are expressed in the exponential phase [30], the challenge posed was more targeted towards DNA sequence patterns that fine-tune gene expression rather than simply determine the ‘on/off’ expression status. RP transcription regulation occurs in a highly coordinated manner and is critical for growth, allowing cells to adjust their protein synthesis capacity to physiological needs [47,48]. This is especially crucial as RP gene expression accounts for 50% of transcripts produced by RNA polymerase II [49] and their dysregulation leads to reduced fitness [47,48]. The yeast genome contains 137 RP genes, of which 19 encode a unique RP and 59 are duplicated. The proper functioning of ribosomes requires that all the ribosome components be expressed in equimolar concentrations [50] while simultaneously remaining responsive to physiological needs [51,52]. This is potentially challenging given the copy-number differences between the RP genes because high copy number genes generally show increased expression. The regulatory mechanisms underlying this fine-tuned regulation are not known. By accurately predicting the activity of the RP genes using the promoter sequences, we demonstrate that a considerable amount of this information is encoded in the DNA sequence.

It is intriguing that our model did not explicitly use transcription factor binding site information and focused only on the 100 bp upstream region. Some of the features identified by our model may influence transcription factor binding or nucleosomes indirectly, and could even affect mRNA translation. Transcription factors are critical for gene regulation. Their empirically identified binding sites are 6 to 8 bp, theoretically putting an upper bound on the level of regulatory flexibility that can be attained by mutating positions at these sites [30,38]. Cooperation between transcription factors or competition among them [15-17], and with nucleosomes [23], provides an additional mechanism for fine-tuned gene expression. RP promoters with high activity have not only more nucleosome disfavoring sequences but also characteristic spatial organization of the binding sites for *Rap1, Sfp1* and *Fhl1* [30]. The low performance of our model on synthetic promoters containing targeted mutations in transcription factor binding sites and nucleosome disfavoring sequences reinforces the importance of these factors. Consistent with this, the combination of our model and the mechanistically driven model involving transcription factors and nucleosome binding [30] was more predictive of promoter activity [34]. Our findings have implications for understanding the fine-tuned regulation of RP genes and engineering desirable activity in synthetic promoters.

## Methods

### DREAM6 Challenge Data

The training data composed of DNA sequence for 90 yeast ribosomal protein (RP) promoters with known activities and a test dataset of 53 promoters was downloaded from the DREAM challenge website (https://www.synapse.org/#!Synapse:syn2820426/wiki/). Details of promoter construction are available from Zeevi et al. 2011 [30] and the DREAM website. Briefly, the organizers considered the promoter region as the sequence 1200 bp upstream of a gene or until the nearest gene. Each promoter was linked to a URA3 selection marker and inserted into the same fixed genomic location of a master yeast strain containing the YFP gene. In total, 110 natural RP promoter strains and 33 strains with synthetically mutated RP promoters were constructed. As a control for experimental variation, all these strains contained a control promoter (TEF2) driving the expression of red fluorescent protein (mCherry). The *mCherry, TEF2, URA3, RP* promoter and *YFP* were all a single contiguous DNA sequence arranged in that order. Measurements of the mCherry expression levels and replicates of promoters had very low variation, enabling the distinction between any two promoters with activities differing by as little as ~8%. The promoter activity was determined as the amount of YFP fluorescence produced during the exponential growth phase, divided by the integral of the OD during the same period. The promoter activity measures the average amount of YFP produced from each promoter, per cell, per second during the exponential phase.

### Feature Extraction

Each promoter sequence was divided into 100 bp non-overlapping windows. The full promoter sequence was considered as another window. To extract information from each of the windows, we considered the frequencies of specific sequences in k-mers (k = 1 to 5), length of homopolymeric stretches DNA mechanical properties (deformability, bendability and stiffness) and nucleosome binding. K-mer counts were performed using custom scripts. DNA mechanical properties were computed using workflows constructed in the Taverna Workbench [53] and BioMoby web-services imported from the Molecular Modeling and Bioinformatics Group, Barcelona, Spain [54]. Bendability was estimated based on trinucleotide parameters obtained from DNase I digestion and nucleosome binding data [35]. Deformability was based on parameters from the analysis of protein-DNA crystallography structures [36]. Bending stiffness was based on bending free energy using the near-neighbor model [37]. Nucleosome binding was based on trinucleotide preferences [55].

### Feature Selection

For each window, feature selection was performed using a linear regression wrapper in the WEKA machine learning toolkit [56] to select feature combinations that are most predictive of promoter activity. Performance of feature combinations was tested using 5- and 10-fold cross-validation.

### Machine Learning Model Exploration

Three models implemented in the WEKA toolkit [56] were considered: support vector machines (SVM) regression using sequential minimal optimization (SMO), linear regression and regression trees. Models were trained using 66% of the data and tested using 34%, and included only the features that were selected as important by the linear regression wrapper. Performance was determined using Pearson correlation between model predictions and actual promoter activities. The SVM model was selected for refinement based on high performance compared to the other models.

### Application of SVM Model to DREAM6 Test Set

Promoter activities were not available to the participants of the challenge. We applied the ensemble of 501 SVMs built from 500 different training/test sets in which 80% of the data was used in training and 20% in testing and a single SVM validated by 66% training set and 34% testing sets. Each SVM model utilized the 24 features selected by a linear regression wrapper as most predictive of promoter activity. To predict activities of the DREAM6 test set, the 24 features were extracted from the upstream 100 bp sequence for each promoter. Predictions were then made using each of the SVM models and averaged to obtain the final predictions.

### Validation of Model by DREAM6 Consortium

Predictions from the SVM ensemble were submitted through the DREAM website to the organizers for a blinded evaluation on the test set. The DREAM organizers used 4 statistics and corresponding *P*-values to evaluate the performance on the test set [34]. Details of the equations used for these statistics have been published separately by the DREAM6 Promoter Prediction Consortium [34].

i. Pearson correlation between predicted and observed activities for each model submitted:To generate *P*–value for observing a Pearson correlation coefficient of the same magnitude or smaller than that of a given participant, a null distribution was generated by randomly sampling predictions from other teams and repeating this 10,000 times [34].
ii. Spearman correlation for participant between ranks of the predicted and actual ranks of promoter activities: A *P*-value was then generated using a null distribution obtained from randomly sampling the predictions made by the other participants. The process was repeated 10,000 times [34].
iii. Chi-square distance metric measuring the distance between predicted and actual promoter activities: To generate a *P*-value for observing a chi-square distance metric of the same magnitude or smaller than that of a given model submission, a null distribution was generated by randomly sampling predictions from other teams and repeating this 10,000 times [34].
iv. A rank distance metric measuring the difference in ranks between predicted ranks and actual ranks of promoter activities. A *P*-value was generated from a null distribution obtained by randomly sampling predicted ranks from other teams, repeating this 10,000 times.

The overall score was defined as the product of the four *P*-values [34].

## Acknowledgements

This work would not have been possible without the pre-publication provision of data to the DREAM challenge by Dr. Eran Segal and his group at the Weizmann Institute of Science, Israel, and the curation of the challenge by the DREAM committee: Drs. Gustavo Stolovitzky, Pablo Meyer and Rachel Norel at IBM Research, USA. We are grateful to the DREAM6 Promoter Prediction Consortium for the rigorous evaluation of the models.

## References

1. Schadt EE, Monks SA, Drake TA, Lusis AJ, Che N, et al. (2003) Genetics of gene expression surveyed in maize, mouse and man. Nature 422: 297–302.

2. Tirosh I, Reikhav S, Sigal N, Assia Y, Barkai N. (2010) Chromatin regulators as capacitors of interspecies variations in gene expression. Mol Syst Biol 6: 435.

3. Tirosh I, Weinberger A, Carmi M, Barkai N. (2006) A genetic signature of interspecies variations in gene expression. Nat Genet 38: 830–834.

4. Field Y, Fondufe-Mittendorf Y, Moore IK, Mieczkowski P, Kaplan N, et al. (2009) Gene expression divergence in yeast is coupled to evolution of DNA-encoded nucleosome organization. Nat Genet 41: 438–445.

5. Gonzales JM, Patel JJ, Ponmee N, Jiang L, Tan A, et al. (2008) Regulatory hotspots in the malaria parasite genome dictate transcriptional variation. PLoS Biol 6: e238.

6. Ellis T, Wang X, Collins JJ. (2009) Diversity-based, model-guided construction of synthetic gene networks with predicted functions. Nat Biotechnol 27: 465–471.

7. Gertz J, Cohen BA. (2009) Environment-specific combinatorial cis-regulation in synthetic promoters. Mol Syst Biol 5: 244.

8. Gertz J, Siggia ED, Cohen BA. (2009) Analysis of combinatorial cis-regulation in synthetic and genomic promoters. Nature 457: 215–218.

9. Kim HD, Shay T, O’Shea EK, Regev A. (2009) Transcriptional regulatory circuits: Predicting numbers from alphabets. Science 325: 429–432.

10. Segal E, Widom J. (2009) From DNA sequence to transcriptional behaviour: A quantitative approach. Nat Rev Genet 10: 443–456.

11. Takahashi K, Yamanaka S. (2006) Induction of pluripotent stem cells from mouse embryonic and adult fibroblast cultures by defined factors. Cell 126: 663–676.

12. Kim HD, O’Shea EK. (2008) A quantitative model of transcription factor-activated gene expression. Nat Struct Mol Biol 15: 1192–1198.

13. Irie T, Park SJ, Yamashita R, Seki M, Yada T, et al. (2011) Predicting promoter activities of primary human DNA sequences. Nucleic Acids Res 39: e75.

14. Cookson W, Liang L, Abecasis G, Moffatt M, Lathrop M. (2009) Mapping complex disease traits with global gene expression. Nat Rev Genet 10: 184–194.

15. Karczewski KJ, Tatonetti NP, Landt SG, Yang X, Slifer T, et al. (2011) Cooperative transcription factor associations discovered using regulatory variation. Proc Natl Acad Sci U S A 108: 13353–13358.

16. Mjolsness E. (2007) On cooperative quasi-equilibrium models of transcriptional regulation. J Bioinform Comput Biol 5: 467–490.

17. Das D, Banerjee N, Zhang MQ. (2004) Interacting models of cooperative gene regulation. Proc Natl Acad Sci U S A 101: 16234–16239.

18. Lam FH, Steger DJ, O’Shea EK. (2008) Chromatin decouples promoter threshold from dynamic range. Nature 453: 246–250.

19. Mirny LA. (2010) Nucleosome-mediated cooperativity between transcription factors. Proc Natl Acad Sci U S A 107: 22534–22539.

20. Li XY, Thomas S, Sabo PJ, Eisen MB, Stamatoyannopoulos JA, et al. (2011) The role of chromatin accessibility in directing the widespread, overlapping patterns of drosophila transcription factor binding. Genome Biol 12: R34–2011–12–4-r34. Epub 2011 Apr 7.

21. Choi JK, Kim YJ. (2009) Intrinsic variability of gene expression encoded in nucleosome positioning sequences. Nat Genet 41: 498–503.

22. Lidor Nili E, Field Y, Lubling Y, Widom J, Oren M, et al. (2010) p53 binds preferentially to genomic regions with high DNA-encoded nucleosome occupancy. Genome Res 20: 1361–1368.

23. Raveh-Sadka T, Levo M, Segal E. (2009) Incorporating nucleosomes into thermodynamic models of transcription regulation. Genome Res 19: 1480–1496.

24. Segal E, Widom J. (2009) Poly(dA:dT) tracts: Major determinants of nucleosome organization. Curr Opin Struct Biol 19: 65–71.

25. Kaplan N, Moore IK, Fondufe-Mittendorf Y, Gossett AJ, Tillo D, et al. (2009) The DNA-encoded nucleosome organization of a eukaryotic genome. Nature 458: 362–366.

26. van der Heijden T, van Vugt JJ, Logie C, van Noort J. (2012) Sequence-based prediction of single nucleosome positioning and genome-wide nucleosome occupancy. Proc Natl Acad Sci U S A 109: E2514–22.

27. Segal E, Widom J. (2009) What controls nucleosome positions? Trends Genet 25: 335–343.

28. Lee CK, Shibata Y, Rao B, Strahl BD, Lieb JD. (2004) Evidence for nucleosome depletion at active regulatory regions genome-wide. Nat Genet 36: 900–905.

29. Shivaswamy S, Bhinge A, Zhao Y, Jones S, Hirst M, et al. (2008) Dynamic remodeling of individual nucleosomes across a eukaryotic genome in response to transcriptional perturbation. PLoS Biol 6: e65.

30. Zeevi D, Sharon E, Lotan-Pompan M, Lubling Y, Shipony Z, et al. (2011) Compensation for differences in gene copy number among yeast ribosomal proteins is encoded within their promoters. Genome Res 21: 2114–2128.

31. Yang YH, Dudoit S, Luu P, Lin DM, Peng V, et al. (2002) Normalization for cDNA microarray data: A robust composite method addressing single and multiple slide systematic variation. Nucleic Acids Res 30: e15.

32. Oshlack A, Wakefield MJ. (2009) Transcript length bias in RNA-seq data confounds systems biology. Biol Direct 4: 14–6150–4–14.

33. Kalir S, McClure J, Pabbaraju K, Southward C, Ronen M, et al. (2001) Ordering genes in a flagella pathway by analysis of expression kinetics from living bacteria. Science 292: 2080–2083.

34. Meyer P, Siwo G, Zeevi D, Sharon E, Norel R, et al. (2013) Inferring gene expression from ribosomal promoter sequences, a crowdsourcing approach. Genome Res.

35. Brukner I, Sanchez R, Suck D, Pongor S. (1995) Trinucleotide models for DNA bending propensity: Comparison of models based on DNaseI digestion and nucleosome packaging data. J Biomol Struct Dyn 13: 309–317.

36. Olson WK, Gorin AA, Lu XJ, Hock LM, Zhurkin VB. (1998) DNA sequence-dependent deformability deduced from protein-DNA crystal complexes. Proc Natl Acad Sci U S A 95: 11163–11168.

37. Sivolob AV, Khrapunov SN. (1995) Translational positioning of nucleosomes on DNA: The role of sequence-dependent isotropic DNA bending stiffness. J Mol Biol 247: 918–931.

38. Raveh-Sadka T, Levo M, Shabi U, Shany B, Keren L, et al. (2012) Manipulating nucleosome disfavoring sequences allows fine-tune regulation of gene expression in yeast. Nat Genet 44: 743–750.

39. Lascaris RF, Mager WH, Planta RJ. (1999) DNA-binding requirements of the yeast protein Rap1p as selected in silico from ribosomal protein gene promoter sequences. Bioinformatics 15: 267–277.

40. Packer MJ, Dauncey MP, Hunter CA. (2000) Sequence-dependent DNA structure: Tetranucleotide conformational maps. J Mol Biol 295: 85–103.

41. Laurens N, Rusling DA, Pernstich C, Brouwer I, Halford SE, et al. (2012) DNA looping by FokI: The impact of twisting and bending rigidity on protein-induced looping dynamics. Nucleic Acids Res 40: 4988–4997.

42. Starr DB, Hoopes BC, Hawley DK. (1995) DNA bending is an important component of site-specific recognition by the TATA binding protein. J Mol Biol 250: 434–446.

43. Vijayan V, Zuzow R, O’Shea EK. (2009) Oscillations in supercoiling drive circadian gene expression in cyanobacteria. Proc Natl Acad Sci U S A 106: 22564–22568.

44. Parvin JD, McCormick RJ, Sharp PA, Fisher DE. (1995) Pre-bending of a promoter sequence enhances affinity for the TATA-binding factor. Nature 373: 724–727.

45. Bosio MC, Negri R, Dieci G. (2011) Promoter architectures in the yeast ribosomal expression program. Transcription 2: 71–77.

46. Yonetani Y, Kono H. (2009) Sequence dependencies of DNA deformability and hydration in the minor groove. Biophys J 97: 1138–1147.

47. Li B, Vilardell J, Warner JR. (1996) An RNA structure involved in feedback regulation of splicing and of translation is critical for biological fitness. Proc Natl Acad Sci U S A 93: 1596–1600.

48. Deutschbauer AM, Jaramillo DF, Proctor M, Kumm J, Hillenmeyer ME, et al. (2005) Mechanisms of haploinsufficiency revealed by genome-wide profiling in yeast. Genetics 169: 1915–1925.

49. Warner JR. (1999) The economics of ribosome biosynthesis in yeast. Trends Biochem Sci 24: 437–440.

50. Spahn CM, Beckmann R, Eswar N, Penczek PA, Sali A, et al. (2001) Structure of the 80S ribosome from saccharomyces cerevisiae--tRNA-ribosome and subunit-subunit interactions. Cell 107: 373–386.

51. Ju Q, Warner JR. (1994) Ribosome synthesis during the growth cycle of saccharomyces cerevisiae. Yeast 10: 151–157.

52. Causton HC, Ren B, Koh SS, Harbison CT, Kanin E, et al. (2001) Remodeling of yeast genome expression in response to environmental changes. Mol Biol Cell 12: 323–337.

53. Oinn T, Addis M, Ferris J, Marvin D, Senger M, et al. (2004) Taverna: A tool for the composition and enactment of bioinformatics workflows. Bioinformatics 20: 3045–3054.

54. Goni JR, Fenollosa C, Perez A, Torrents D, Orozco M. (2008) DNAlive: A tool for the physical analysis of DNA at the genomic scale. Bioinformatics 24: 1731–1732.

55. Satchwell SC, Drew HR, Travers AA. (1986) Sequence periodicities in chicken nucleosome core DNA. J Mol Biol 191: 659–675.

56. Hall M, Frank E, Holmes G, Pfahringer B, Reutemann P, et al. (2009) The WEKA data mining software: An update. SIGKDD Explorations, 11.

